# Viral receptor profiles of masked palm civet revealed by single-cell transcriptomics

**DOI:** 10.1101/2021.07.20.452903

**Authors:** Dongsheng Chen, Zhihua Ou, Haoyu Wang, Yanan Zhang, Jiacheng Zhu, Fuyu An, Jinqian Xu, Xiangning Ding, Peiwen Ding, Lihua Luo, Weiying Wu, Qiuyu Qin, Yanan Wei, Wandong Zhao, Zhiyuan Lv, Tianming Lan, Meiling Li, Wensheng Zhang, Huan Liu, Yan Hua

**Affiliations:** BGI-Shenzhen, Shenzhen 518083, China; Shenzhen Key Laboratory of Unknown Pathogen Identification, BGI-Shenzhen, Shenzhen 518083, China; College of Life Sciences, University of Chinese Academy of Sciences, Beijing 100049, China; Tsinghua-Berkeley Shenzhen Institute, Tsinghua University, Shenzhen, China; Guangdong Provincial Key Laboratory of Silviculture, Protection and Utilization, Guangdong Academy of Forestry, Guangzhou 510520, China; School of Basic Medicine, Qingdao University, Qingdao 266071, China; School of Basic Medical Sciences, Binzhou Medical University, No. 346, Guanhai Road, Laishan District, Yantai City, Shandong, China

## Abstract

Civets are small mammals belonging to the family *Viverridae*. The masked palm civets (*Paguma larvata*) served as an intermediate host in the bat-to-human transmission of severe acute respiratory syndrome coronavirus (SARS-CoV) in 2003^1^. Because of their unique role in the SARS outbreak, civets were suspected as a potential intermediate host of SARS-CoV-2, the etiological pathogen of the COVID-19 pandemic. Besides their susceptibility to coronaviruses, civets can also be infected by other viruses, such as canine distemper viruses^2^, parvoviruses^3^, influenza viruses^4^, etc. Regarding the ecological and economical role of civets, it is vital to evaluate the potential threats from different pathogens to these animals. Receptor binding is a necessary step for virus entry into host cells. Understanding the distribution of receptors of various viruses provides hints to their potential tissue tropisms. Herein, we characterized the cell atlas of five important organs (the frontal lobe, lung, liver, spleen and kidney) of masked palm civets (*Paguma larvata*) and described the expression profiles of receptor associated genes of 132 viruses from 25 families, including 16 viruses from 10 families reported before that can attack civets and 116 viruses with little infection record.

## Results

To build a comprehensive cell atlas of civet organs, we performed single-cell RNA sequencing to five organs of an adult male masked palm civet. After pre-processing, a total of 66,553 cells (Fig. 1a), including 6,593 cells from the frontal lobe, 13,009 cells from the lung, 34,883 cells from the liver, 10,138 cells from the spleen and 1,930 cells from the kidney were acquired. We conducted unsupervised clustering and annotated resulted cell clusters based on canonical markers (Fig. 1b-f, Table S1). In the frontal lobe, we identified 8 major cell types for 25 clusters, including astrocytes (AST), excitatory neurons (EX), inhibitory neurons (IN), oligodendrocytes (OLG), OLG progenitor cells (OPC), microglia (MG), smooth muscle cells (SMC), and endothelial cells (END). In lung, 7 cell types were annotated for 30 clusters, including pulmonary alveolar type I (AT1), pulmonary alveolar type II (AT2), macrophages (MAC), epithelial cells (EC), ciliated cells (CC), fibroblasts (FIB) and END. 5 cell types were characterized for 19 clusters in liver, including hepatic stellate cells (HSC), immune cells (IC), liver sinusoidal endothelial cells (LSEC), cholangiocytes and hepatocytes. In spleen, we annotated 21 clusters to 11 cell types, including END, FIB, neural cells (NEU), and several immune cell types, which were naive T cells, T cells, T follicular helper cells (Tfh), regulatory T cells (Treg), B cells, macrophages (MAC), dendritic cells (DC) and natural killer cells (NK). In kidney, 11 clusters were characterized to 8 cell types, including proximal tubule cells (PCT), distal convoluted tubule cells (DCT), podocytes (Podo), loop of Henle cells (LOH), collecting duct intercalated cells (CD-IC), collecting duct principal cells (CD-PC), pericytes (PER) and END. To share the single cell atlas of civet, we constructed an online platform, http://120.79.46.200:81/Civet, allowing researchers freely exploring our data set and analysis results.

**Fig. 1.**
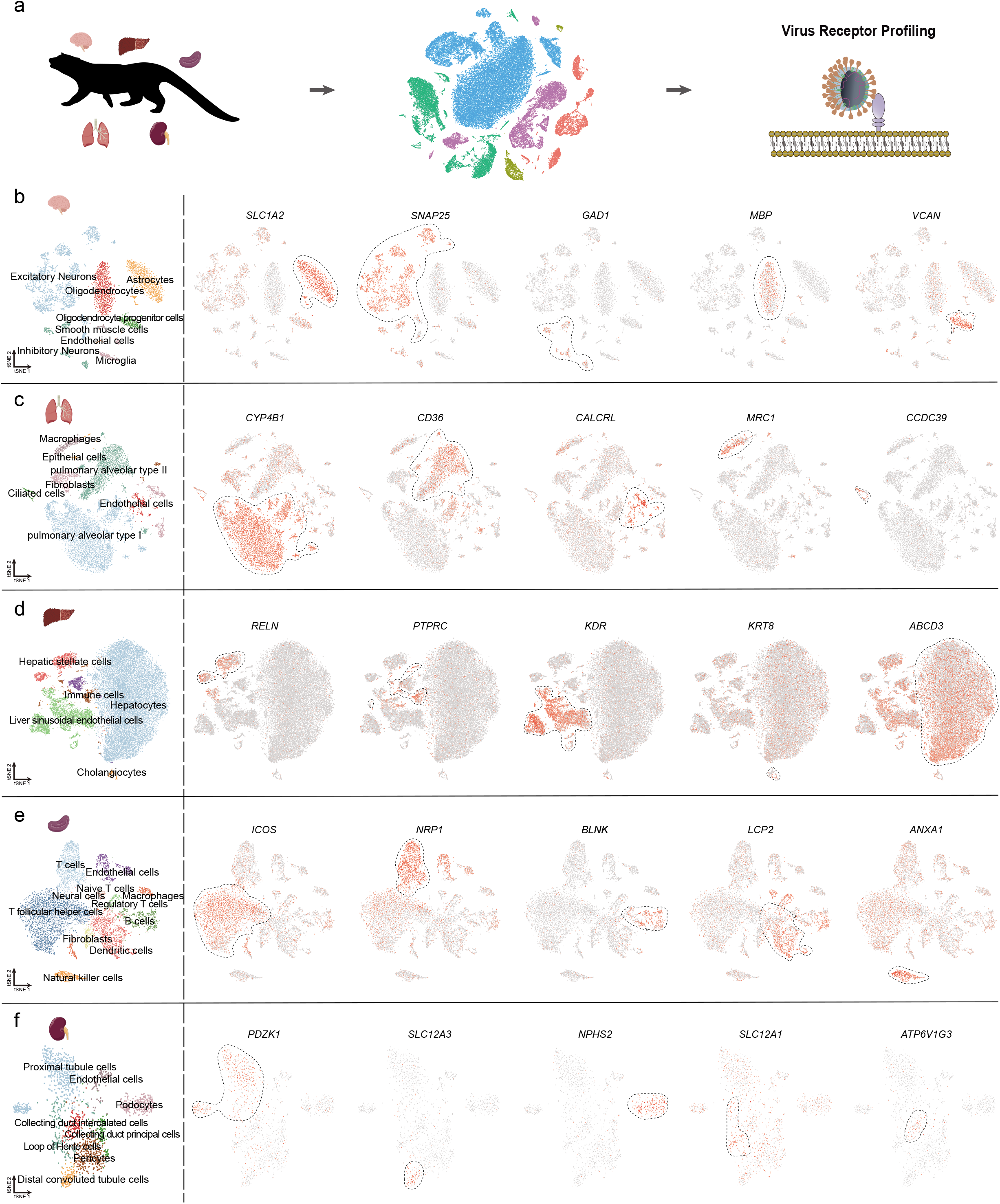
Single cell atlas of the frontal lobe, lung, liver, spleen and kidney of civet. **a** Workflow of this study. **b** tSNE plot of the frontal lobe cells. Colors represent different cell types. Feature plots indicate the expression of cell markers with red color indicating high expression patterns. **c** tSNE plot of lung cells and feature plots of cell markers. **d** tSNE plot of liver cells and feature plots of cell markers. **e** tSNE plot of spleen cells and feature plots of cell markers. **f** tSNE plot of kidney cells and feature plots of cell markers.

Among the viruses in analysis, we placed special focus on six virus families that prone to cause inter-species transmissions associated with severe diseases, including *Coronaviridae, Filoviridae, Orthomyxoviridae, Paramyxoviridae, Parvoviridae* and *Rhabdoviridae*, though no civet infections by filoviruses have yet been reported. For *Coronaviridae, ACE2*, the common receptor of SARS-CoV, SARS-CoV-2 and human coronavirus NL63, was expressed by a small fraction of cells in the frontal lobe, lung, liver, spleen and kidney, with the highest expression level in loop of Henle cells and proximal tubule cells of the kidney. *AXL*, the common receptor of SARS-CoV-2, ebolavirus and marburg marburgvirus, showed the same expression pattern as *ACE2*. Importantly, NRP1, another receptor for SARS-CoV-2, was widely detected in all five organs and expressed in a much higher level than ACE2. *ANPEP*, a common receptor for animal and human multiple coronaviruses, was only expressed by a small fraction of hepatocytes. For *Filoviridae*, multiple receptors (*ITCH, NPC1* and *MERTK*) of Ebola viruses and Marburg viruses displayed high expression in the five organs, though the expression levels varied among cell types. For *Orthomyxoviridae*, three receptor-associated genes for avian influenza A virus (*UVRAG, ANXA5* and *EGFR*) and one receptor-associated gene for bat influenza A virus H18N11 (*CD74*) showed high expression patterns in all five organs, especially in the lung and spleen. For *Paramyxoviridae, SLAMF1* and *NECTIN4* were both receptors of canine morbilliviruses but *SLAMF1* was detected in a small number of cells across the five organs while *NECTIN4* was only highly expressed by a few lung cells. *EFNB2* is a receptor of Henipaviruses, which was detected in all five organs with high expressions in the lung, spleen and kidney. For *Parvoviridae, TFRC*, which is a common receptor of parvoviruses, was widely expressed in the five organs with a significant enrichment in the endothelial cells of the frontal lobe. For *Rhabdoviridae*, the receptors of lyssaviruses, *GRM2* and *NCAM1*, were detected in all five organs with the highest expression in the frontal lobe. The receptors of vesicular stomatitis virus, *UVRAG* and *LDLR*, were identified in five organs with lung cells showing the highest expressions (Fig. S1).

Besides the above six families, we also categorized the receptor distributions of other viruses capable of infecting civets. *F11R*, the receptor for viruses of *Caliciviridae* and *Reoviridae*, was only detected in the civet lung (AT1 and AT2) and liver (liver sinusoidal endothelial cells and hepatocytes). Another two receptors of reoviruses, *RTN4R* and *ITGB1*, were both widely expressed in all five organs. *CXCR4*, a receptor of feline immunodeficiency virus (*Retroviridae*), was widely detected in the frontal lobe, lung, liver and spleen at low levels. *TLR8* and *TLR7*, the receptors of severe fever with thrombocytopenia syndrome virus (*Bunyavirales, Phenuiviridae*), were both found in the lung and spleen but *TLR8* was also distributed in the frontal lobe and liver. The receptor of West Nile virus (*Flaviviridae*), *ITGAV*, showed expression in all five organs (Fig. S2).

The receptor expressions of other viruses that have not been reported to infect civets were also identified, including *Adenoviridae, Arenaviridae, Flaviviridae, Hepadnaviridae, Herpesviridae, etc*. Multiple receptors for *Adenoviridae* (*WWP2, CXADR, ITGB5 and ITGAV*), *Herpesviridae* (*ITGB1, CR1, ITCH, ITGB8, ITGAV* and *IDE*), *Picornaviridae* (*ITGAV, CXADR* and *ITGB8*) and *Reoviridae* (*ITGB1, HSPA8* and *ITGAV*) were abundantly expressed by cells of the five organs. Expressions of the other receptors tended to be lower than the above-mentioned ones or concentrated in certain organs (Fig. S1, Fig. S2).

The receptor expression profiles could help us better clarify or predict the pathological outcomes caused by viral infection in civets. For example, the canine parvoviruses were reported to cause diarrhea and deaths in civets and the viral DNA can be detected in the brain, liver, heart, spleen and small intestine^3^, which is consistent with the wide distribution of its receptor *TFRC* in the civet organs. Lyssaviruses usually cause fatal encephalitic diseases in a wide range of mammals with civet infections reported in Africa and Asia^5,6^. Our results showed an obvious enrichment of a related receptor, *NCAM1*, in multiple cell types of the civet frontal lobe, which may contribute to the neurovirulence of these viruses. The receptor of SARS-CoV-2, ACE2, was only expressed at moderate levels in the kidney and flow cytometric experiment showed undetectable binding between the civet ACE2 ortholog and the viral receptor-binding domain. However, two alternative receptors, NRP1 and AXL, were both highly expressed in the lung and spleen, indicating potential susceptibility of civets to SARS-CoV-2, although *in vivo* infection remains unclear.

## Discussion

Taken together, we have built a comprehensive multi-organ cell atlas of masked palm civet and described the distribution of various viral receptors in these tissues, providing preliminary evidence of the potential tissue tropism of these viruses in civets. The results could enhance our knowledge of the biological background of civets and their susceptibility to various pathogens, which may facilitate the control and prevention of enzootic and zoonotic viruses.

## Acknowledgement

This work was supported by China National GeneBank (CNGB).

## AUTHOR CONTRIBUTIONS

Y.H., H. L., D.C. and Z. O. conceived and designed the project. J.Z., F.A., J.X., were responsible for sample collection and dissection. W.W. participated in single-nucleus library construction and sequencing. H.W. performed single cell analysis. Y.Z., H.W., Z. O., D.C., X.D., P.D., L.L., Q.Q., Y.W., W.D., Z.L., T.L., M.L., W.Z., participated in data interpretation, data visualization and manuscript writing. Y.H., H. L. revised the manuscript.

## Conflict of Interest

The authors declare no competing interests.

## Supplementary materials

### Supplementary Figures

**Fig. S1. Distribution of viral receptors for *Coronaviridae, Filoviridae, Orthomyxoviridae, Paramyxoviridae, Parvoviridae* and *Rhabdoviridae* in the frontal lobe, lung, liver, spleen and kidney of civet.** Bubble plot shows the viral receptor expressions in different organs. Dot size represents the percentage of cells expressing the corresponding receptor. Color saturation indicates the average scaled expression level. Viruses that are capable of infecting civets are in red text.

**Fig. S2. Distribution of other viral receptors in civet organs.** Bubble plot shows the expressions of viral receptors in different organs. Viruses that are capable of infecting civets are in red text.

### Supplementary Tables

Table S1. Marker genes for cell types of frontal lobe, lung, liver, spleen and kidney

Table S2. Expression of viral receptor genes in civet

Table S3. DEG of cell types in frontal lobe, lung, liver, spleen and kidney

Table S4. GO term of cell type DEGs in frontal lobe, lung, liver, spleen and kidney

## Notes

### Competing Interest Statement

The authors have declared no competing interest.

## Reference

1. Guan, Y. et al. Isolation and characterization of viruses related to the SARS coronavirus from animals in Southern China. Science (80-.). 302, 276–278 (2003).

2. Techangamsuwan, S. et al. Pathologic and Molecular Virologic Characterization of a Canine Distemper Outbreak in Farmed Civets. Vet. Pathol. 52, 724–731 (2015).

3. Mendenhall, I. H. et al. Evidence of canine parvovirus transmission to a civet cat (Paradoxurus musangus) in Singapore. One Heal. 2, 122–125 (2016).

4. Roberton, S. I. et al. Avian influenza H5N1 in viverrids: Implications for wildlife health and conservation. Proc. R. Soc. B Biol. Sci. 273, 1729–1732 (2006).

5. Sabeta, C. T. et al. Rabies in the African civet: An incidental host for lyssaviruses? Viruses 12, (2020).

6. Matsumoto, T. et al. Novel sylvatic rabies virus variant in endangered golden palm civet, Sri Lanka. Emerg. Infect. Dis. 17, 2346–2349 (2011).

