## Supplementary Information for "Viral receptor profiles of masked palm civet revealed by single-cell transcriptomics"

**Material and Methods**

**Ethics statement**

This project was approved by the Institutional Review Board on Ethics Committee of BGI (Approval letter reference number BGI-NO. BGI-IRB A20008). All procedures were conducted according to the guidelines of the Institutional Review Board on the Ethic Committee of BGI. The ‘Guidelines on Ethical Treatment of Experimental Animals’ established by the Ministry of Science and Technology, China was strictly followed for sampling procedures.

**Sample collection and nuclei extraction**

The frontal lobe, lung, liver, spleen and kidney were collected from an adult male civet. The collected organs were washed with 1X phosphate buffered saline (PBS), then quickly frozen and stored in liquid nitrogen. The mechanical extraction method^1^ was used for nuclei extraction and separation.

**Single nuclei library construction and sequencing**

Single nucleus libraries of frontal lobe, lung, liver, spleen and kidney of civet were constructed using Single Cell 3ʹ GEM, Library & Gel Bead Kit v3 (PN-1000075) following the standard manufacturer’s instructions. Library conversion was performed using MGIEasy Universal DNA Library Preparation Reagent Kit to compatible with BGISEQ-500 sequencing platform.

**Single-nucleus RNA-sequencing data processing**

Single-nucleus RNA-sequencing (snRNA-seq) data and gene expression matrix were obtained using Cell Ranger 3.0.2 (10X Genomics). Raw data was filtered using package Seurat before downstream analysis, and cells with nFeature_RNA > 200 and percent.mt < 10 were retained. The R package Seurat v3^2^ was used for the data normalization and highly variable genes (HVGs) recognition. The principal component analysis (PCA) was performed using HGVs and the principal components (PCs) significance was calculated for cell cluster identification and visualization. “RunTSNE” function was applied for cluster visualization. Double cells were defined and removed using DoubletFinder package.

**Differential expression analysis, functional enrichment analysis and cell types annotation**

Differentially expressed genes (DEGs) were identified by “FindAllMarkers” function in Seurat. For DEGs of each cluster, we applied clusterProfiler Package for GO enrichment analysis. *P*-values were adjusted with Benjaminiand Hochberg (BH) method and pathways with an adjust p-values ≤ 0.05 were considered significant enriched DEGs, GO pathways and verified markers were combined to annotate cell types for each organ.

**Expression of different virus receptors**

All receptors of virus were collected from viral receptor database^3^, Human Lung Cell Atlas (HLCA) database, or published articles. The expression of receptors in all cell types were displayed in dot plot using R package ggplot2.

**Data availability**

The single cell atlas of mink is available via http://120.79.46.200:81/Civet. The raw data supporting the findings of this study will be made available upon request.

**Reference**

1.	Zhu, J. *et al.* Single-cell atlas of domestic pig cerebral cortex and hypothalamus. *Sci. Bull.* (2021) doi:10.1016/j.scib.2021.04.002.

2.	Stuart, T. *et al.* Comprehensive Integration of Single-Cell Data. *Cell* **177**, 1888-1902.e21 (2019).

3.	Zhang, Z. *et al.* Cell membrane proteins with high N-glycosylation, high expression and multiple interaction partners are preferred by mammalian viruses as receptors. *Bioinformatics* **35**, 723–728 (2019).
