## Supplementary figures and images for "Viral receptor profiles of masked palm civet revealed by single-cell transcriptomics"

### Fig. S1

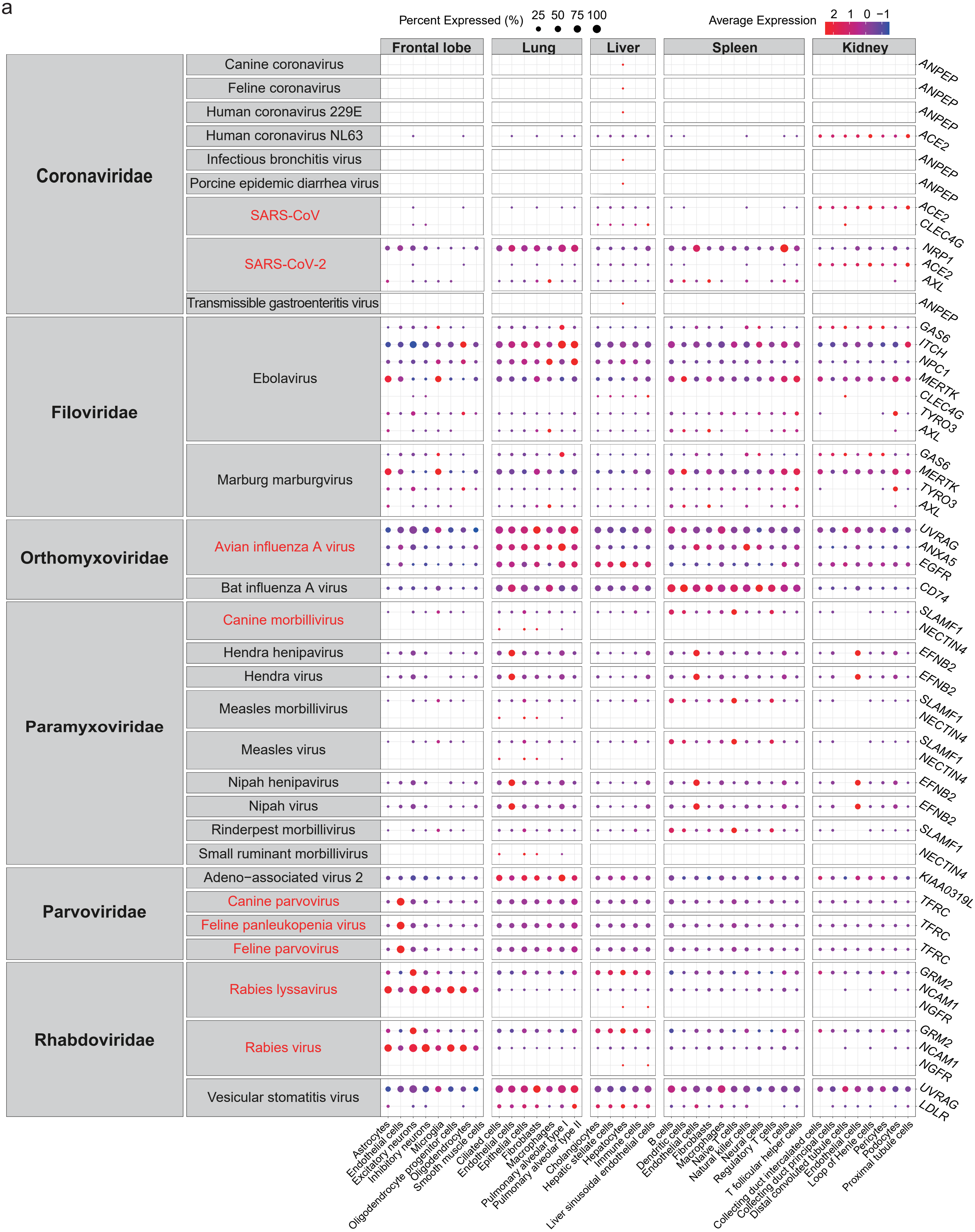

### Fig. S2

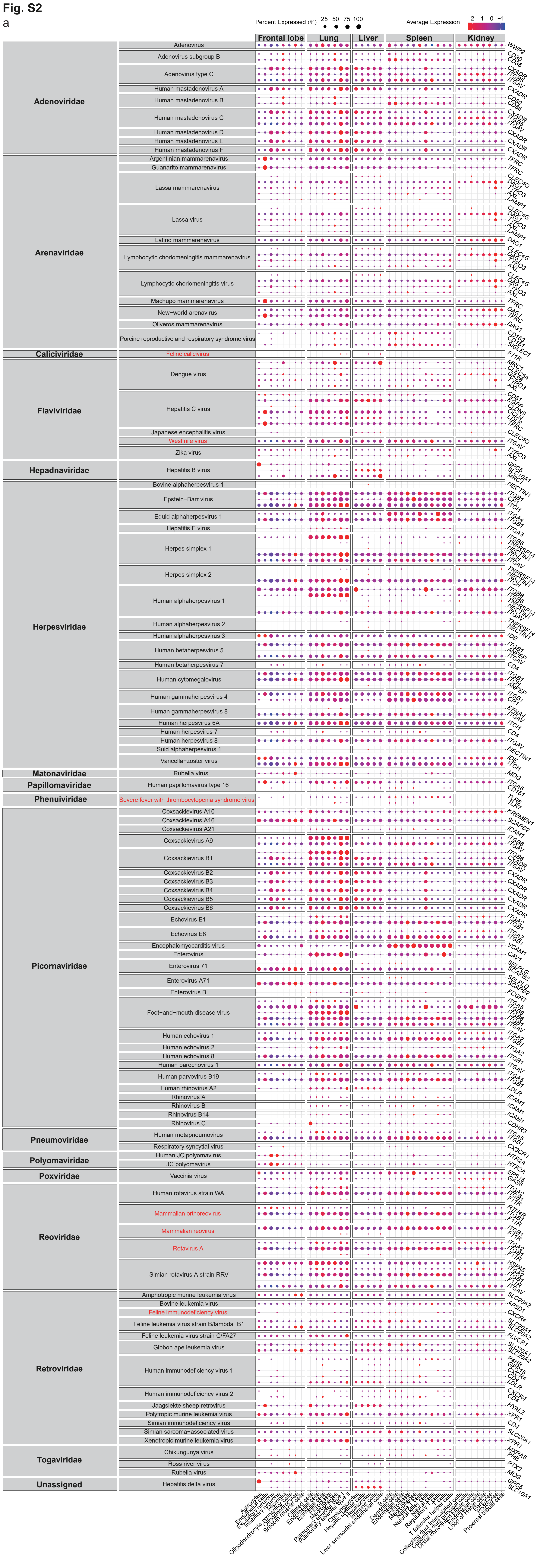
